# Calling the Amino Acid Sequence of a Protein/Peptide from the Nanospectrum Produced by a Sub-nanometer Diameter Pore

**DOI:** 10.1101/2021.10.17.464717

**Authors:** Xiaowen Liu, Zhuxin Dong, Gregory Timp

## Abstract

The blockade current that develops when a protein translocates across a thin membrane through a sub-nanometer diameter pore (i.e., a nanospectrum) informs with extreme sensitivity on the sequence of amino acids that constitute the protein. Whereas mass spectrometry (MS) is still the dominant technology for protein identification, it suffers limitations. In proteome-wide studies, MS fails to sequence proteins *de novo*, but merely classifies a protein and it is not very sensitive requiring about a femtomole to do that. Compared with MS, a sub-nanometer diameter pore (i.e. a sub-nanopore) directly reads the amino acids constituting a single protein molecule, but efficient computational tools are still required for processing and interpreting the blockade current. Here, we delineate computational methods for processing sub-nanopore nanospectra and predicting electrical blockade currents from protein sequences, which are essential for protein identification.

## 1. Introduction

Sequencing proteins by measuring the blockade current through a sub-nanometer diameter pore (sub-nanopore) is a potentially disruptive technology [1–3]. So far, this technology has been successfully employed to analyze histones and other proteins [4, 5]. A sub-nanopore that is sputtered through a nanometer-thick membrane with a tightly focused high-energy electron beam, is designed to be about the size of an <u>a</u>mino <u>a</u>cid (AA), which accounts for the extreme sensitivity. When the membrane is immersed in electrolyte and a voltage is applied across it, the electric force on the ions in solution produces a current through the sub-nanopore. Subsequently, when a charged denatured protein is impelled by the same electric force through the sub-nanopore, the open pore ionic flow is blocked by the acids in the pore waist. The resulting blockade current or nanospectrum is modulated by the AA sequence constituting the protein. It has been shown that the nanospectrum is correlated with the volume of amino acids occluding the pore, so the AA sequence constituting the protein can be read from the fluctuations in the blockade current [1, 3].

Currently, <u>m</u>ass <u>s</u>pectrometry (MS) is the leading technology for protein identification [6]. Whereas bottom-up MS analyzes proteolytically digested short peptides, top-down MS is capable of analyzing intact proteins [7]. However, MS-based protein identification has fundamental limitations in sensitivity and measurable molecular masses [8]. MS detection requires between an attomole and femtomole of protein, making it challenging to identify low abundance peptides or proteins. Moreover, MS often fails to achieve high sequence coverage for long proteins. Bottom-up MS identifies some peptides of long proteins but does not offer high sequence coverage. Topdown MS provides whole sequence coverage of proteins, but the measurable mass of a protein is limited due to mass spectrometers’ capacity.

Sequencing protein with a sub-nanopore could be a disruptive technology for several reasons [2]. First, a sub-nanopore reads single protein molecules, significantly increases the dynamic range of protein identification. Because of this, single-molecule protein sequencing has many applications in low abundance protein analysis and single-cell proteomics. Second, sub-nanopore sequencing is not limited by the molecular weight of the protein. In principle, a sub-nanopore could read thousands of AAs in a single molecule. Third, a sub-nanopore is capable of analyzing the prevalence of heterogeneity in mRNA translation [9] and <u>p</u>ost-<u>t</u>ranslational <u>m</u>odifications (PTMs) [10] by *direct* protein-level analysis. So far, the analysis of blockade currents in nanometer-diameter pores has been applied successfully to call the sequence of bases constituting DNA and RNA. Many methods have been developed for improving the base calling accuracy of nanopore DNA reads, including hidden Markov and neural network models [11]. As a result, the basecalling accuracy has been improved from 63% to >95% within the last several years [12–14]. Similarly, computational methods have the potential to improve the accuracy of sub-nanopore protein sequencing. However, detecting the acid sequence in a protein with a sub-nanopore is more exacting than discriminating the four bases that constitute DNA with a nanopore, and efficient tools for the analysis of the blockade current produced when a protein translocates across a membrane through a sub-nanopore protein are lacking.

The interpretation of sub-nanopore nanospectra, however, still presents a daunting challenge for detection and identification. Current blockade signals are determined mainly by the volume of the AA in the pore. Moreover, there are large variances in the measured blockades that may be due to other factors, which include electrical and molecular configurational noise, the AA mobility and hydrophobicity in the pore, and the neighboring acids in the sequence. Even if it is detected, calling the acid is confounded by the primary structure of a protein, which is drawn from twenty proteinogenic AAs. Beyond just the twenty proteinogenic AAs, the challenge confronting direct protein sequencing is compounded by protein isoforms derived from closely related duplicate genes or the same gene by alternative splicing, proteolytic cleavage, somatic recombination, or PTMs [10, 15]. In a groundbreaking effort, Kolmogorov *et al*. first tackled the analysis by benchmarking several machine learning models and presented an alignment algorithm for protein identification using only a few nanospectra [16], but due to the noise it remained problematic to identify a protein by searching nanospectra against a protein database the size of the human proteome.

The sub-nanopore technology has advanced rapidly in the past several years; it is now capable of measuring the volumes of single amino acids instead of several consecutive amino acids [17]. So, with the proper computational tools, it should be possible to decode single amino acids directly using nanospectra. Here, several computational methods for processing nanospectra and predicting theoretical nanospectra from protein sequences are described. These methods promise to improve the accuracy of theoretical nanospectral prediction and increase the <u>P</u>earson <u>c</u>orrelation <u>c</u>oefficient (PCC) between the empirical and theoretical nanospectra to > 0.9.

## 2. Methods

### 2.1 Peptide synthesis

Two carrier-free peptides were used in experiments (Anaspec, Fremont, CA): amyloid beta 42 (Aβ_1-42_) (DAEFRHDSGYEVHHQKLVFFAEDVGSNKGAIIGLMVGGVVIA), and a scrambled variant with the same chemical constituency as amyloid beta 42 (SAβ_1-42_) (AIAEGDSHVLKEGAYMEIFDVQGHVFGGKIFRVVDLGSHNVA). The two peptides were reconstituted according to the protocols offered by the manufacturer. Typically, the peptides were reconstituted at high (100 μg/ml) concentration in <u>p</u>hosphate-<u>b</u>uffered <u>s</u>aline (1× PBS) without adding <u>b</u>ovine <u>s</u>erum <u>a</u>lbumin (BSA) to avoid false readings. From this solution, aliquots diluted to 2× the concentration of denaturant with 50 pM protein, 20-100 μM beta-mercaptoethanol (BME), 250 mM *NaCl* with 2-5×10^-3^ % <u>s</u>odium <u>d</u>odecyl <u>s</u>ulfate (SDS) were vortexed and heated to 85 °C for 120 min. The solution was allowed to cool (to 5° C) and added in 1:1 proportion with the (75 μL) electrolyte in the reservoir of the polydimethylsiloxane (PDMS) microfluidic device bound to the silicon chip supporting the membrane with a pore through it housed in a 5°C cold room.

### 2.2 Sub-nanopore fabrication and visualization

Custom-made amorphous silicon (a-*Si*) membranes (SiMPore, Inc. West Henrietta, NY) nominally 5 nm thick were manufactured by the method described in [17]. Briefly, amorphous silicon was sputter-deposited on a 50 nm thick thermal *SiO*_2_ layer grown on a float-zone silicon handle 100 μm thick and subsequently capped with another *SiO*_2_ layer with the same 25-50 nm thickness followed by the deposition of 150 nm of tetraethyl orthosilicate (TEOS). A membrane < 4-5 μm on-edge was revealed by an ethylene diamine and pyrocatechol chemical etch of the silicon through a silicon nitride window defined by photolithography on the polished back-side of the handle wafer. Finally, a <u>b</u>uffered <u>o</u>xide etch (10:1 BOE) was used to remove the oxide to produce an a-*Si* membrane, which ranged from *t* = 3.5 to 6 nm thick.

Just prior to loading it into the <u>t</u>ransmission <u>e</u>lectron <u>m</u>icroscopy (TEM) column, the membranes were plasma cleaned using Tergeo-EM (PIE Scientific, Union City, CA USA). The Tergeo-EM was operated at 10 W using an 80% Ar+20% O_2_ gas feed in a down-stream, pulse mode (1/16 duty-cycle, which was cycled twice for a total exposure of 2 min) such that the samples were actually outside the plasma (to eliminate sputtering) and subjected to only extremely short plasma pulses (to reduce the intensity). Subsequently, a pore was sputtered through the thin a-*Si* membrane using a tightly focused, high-energy (300 kV) electron beam carrying a current ranging from 300-800 pA (post-alignment) in a <u>S</u>canning <u>T</u>ransmission <u>E</u>lectron <u>M</u>icroscope (STEM, FEI Titan 80-300 or FEI Themis Z, Hillsboro, OR) with a <u>F</u>ield <u>E</u>mission <u>G</u>un (FEG).

After sputtering, the pore was re-acquired with either <u>H</u>igh-<u>R</u>esolution <u>T</u>ransmission <u>E</u>lectron <u>M</u>icroscopy (HRTEM) or <u>H</u>igh-<u>A</u>ngle <u>A</u>nnular <u>D</u>ark <u>F</u>ield (HAADF-)STEM. To minimize beam damage, the pores were examined using low beam current (<10-30 pA) or low energy (80kV) or both. The illumination convergence angle in the Titan was typically α = 10 mrad at 300kV, whereas in the Themis Z, α = 18 mrad at 300kV or α =27.1 mrad at 80kV with a monochromator limiting the energy dispersion in the range 200-220mV at 80kV according to EELS.

### 2.3 Microfluidics

The silicon chip supporting the membrane with a single pore through it with or without a polyimide laminate was bonded to a polydimethylsiloxane (PDMS, Sylgard 184, Dow Corning) microfluidic device formed using a mold-casting technique [17]. The microfluidic device consisted of two microchannels (each 250 × 75 μm^2^ in cross-section) connected by a *via* that could be as small as 25 μm in diameter. A tight seal was formed between the silicon chip containing the a-*Si* membrane with the pore in it and the PDMS *trans*-side of the microfluidic channel with a plasma-bonding process (PDS-001, Harrick Plasma, Ithaca, NY). Subsequently, two separate *Ag/AgCl* electrodes (Warner Instruments, Hamden, CT) were embedded in each channel to independently, electrically address the *cis*- and *trans*-sides of the membrane. Likewise, the two microfluidic channels were also connected to external pressure and fluid reservoirs through polyethylene tubing at the input and output ports. The port on the *cis*-side was used to convey proteins to the pore.

### 2.4 Low-noise electrical measurements

To perform blockade current measurements, first, the sub-nanopore was wetted by immersion in de-gassed 250 mM *NaCl* electrolyte for 1-3 days [17]. Subsequently, to measure the blockade current, a transmembrane voltage bias (< 700 mV) was applied to the reservoir (containing 75 μL of electrolytic solution and 75 μL of 2× concentrated solution of protein and denaturant) relative to the ground in the channel using *Ag/AgCl* electrodes and the corresponding pore current was measured at 5 ± 0.1°C using either an Axopatch 700B or an Axopatch 200B amplifier with an open bandwidth. The actual bandwidth was inferred from the rise-time to a sharp (10 ps rise-time) input pulse to be about 75 kHz to 100 kHz, depending on the amplifier and the feedback. The analog data were digitized by a 16-bit DigiData 1550B data acquisition system (DAQ, Molecular Devices, Sunnyvale, CA) at a sampling rate of 500 kS/s and recorded in 3 minute-long acquisition windows. Generally, no blockades were observed beyond the noise in controls that comprised the electrolyte and the denaturants (SDS and BME), which were heated to 85°C and then cooled without protein. A total of 12 Axon binary files (ABF) were collected for Aβ_1-42_, and 70 ABF files for SAβ_1-42_.

### 2.5 Data pre-processing

The current blockade signals (nanospectra) in ABF files were extracted using a homemade software package based on OpenNanopore (version 1.2) [18]. Nanospectra with a relatively long duration provided useful information for AA sequencing, but those that are too short did not. So, the nanospectra with a duration shorter than 170 μs were ignored. The duration for a peptide in the sub-nanopore ranged from tens of microseconds to tens of milliseconds, and the numbers of data points in nanospectra vary dramatically. To address the variation in blockade duration, it was assumed that each raw blockade represented the same pattern of fluctuations and so it was converted into a nanospectrum of 500 data points by averaging or interpolating between neighboring data points. Thus, a consensus formed from these spectra represents signals irregularly (nonuniformly) sampled above, at, and below the Nyquest rate. Regardless of the duration, consensuses formed this way can inform on each AA in the sequence [3, 19–23]

### 2.6 Features for AAs

Linear regression was used to predict the current blockade signals of AAs in peptides. Several encoding methods were used for representing amino acids. A given peptide sequence *a*_1_,*a*_2_,…,*a_n_* were converted to a list of AA volumes: *b*_0_,*b*_1_,…, *b*_*n*+1_, where *b*_0_ = *b*_*n*+1_ = 0 and *b_i_* is the AA volume [24] corresponding to *a_i_* for 1 ≤ *i* ≤ *n*. The first encoding method is based on single AA volumes: an AA *a_i_*, is represented by its volume *b_i_*. The second encoding method is based on the volumes of the AA and its two neighboring ones: an AA *a_i_* is represented by two values *b_i_* and *b*_*i*−1_ + *b*_*i*+1_. In the third encoding method, the 20 AAs are divided into 4 groups based on their volumes: minuscule (G, A, S, C), small (T, D, P, N, V), intermediate (E, Q, H, L, I, M, K), and large (R, F, Y, W) [16]. So, given a peptide *a*_1_,*a*_2_,…,*a_n_*, let *M_i_* = 1 if *a_i_* is a minuscule AA and *M_i_* = 0 otherwise, for 1 ≤ *i* ≤ *n*. Specifically, *M*_0_ = *M*_*n*+1_ = 0. For position *i* in the peptide, we extract four features based on the volume of *a_i_*. The first feature *x_M_* is the volume of the AA if it is a minuscule one, and 0 otherwise, defined as *x_M_* = *M_i_b_i_*. The features for small (*x_s_*), intermediate (*x_I_*), and large (*x_L_*) AAs are defined similarly. The three encoding methods are referred to as single AA volume (1AAV), three AA volume (3AAV), and AA group (AAG) methods, respectively.

The three encoding methods were further extended to include a position feature, which represents the distance between the AA and the N- or C-terminus. When the distance is larger than 4, the AA is treated as a middle one and the feature is set to 5. For *a_i_* with position *i*, the position feature *x_P_* is:

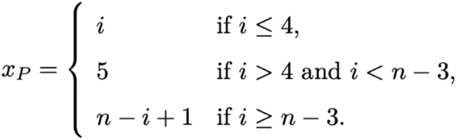

The three encoding methods with the position feature are referred to as 1AAV-P, 3AAV-P, and AAG-P, respectively.

### 2.7 Orientation of the nanospectra

It was assumed that a nanospectrum of a peptide had two possible orientations: a *forward* nanospectrum enters the pore axis N-terminus first and a *backward* nanospectrum C-terminus first. Let *S* = *s*_1_*s*_2_…*s_m_* be an empirical nanospectrum with *m* data points, where *s_i_* is the current blockade signal at time point *i*, and *S*′ = *s_m_s_m−1_*…*s*_1_ the flipped nanospectrum of *S*. To account for the two orientations, a theoretical nanospectrum *T* = *t*_1_*t*_2_… *t_m_* of the peptide derived from the 1AAV model and linear interpolation was generated and compared with empirical nanospectra. *PCC*(*S,T*) represents the PCC of an empirical nanospectrum *S* and the corresponding theoretical nanospectrum *T*. If PCC(*S, T*) > PCC(*S′, T*), then *S* is forward, otherwise, backward. The backward nanospectra were flipped so that all nanospectra have the same orientation.

### 2.8 Dynamic time warping

Let *S* = *s*_1_*s*_2_ …*s_m_* and *T* = *t*_1_*t*_2_ …*t_m_* be an empirical and a theoretical nanospectra of a peptide, respectively. Both *S* and *T* were normalized to have zero mean and unit variance. Let *S*[*i, j*] represent the subsequence *s_i_s_i+1_*… *s_j_* of *S*. Because the velocity of the AAs moving through the sub-nanopore might vary, *s_i_* and *t_i_* might correspond to different AAs in the peptide. To address the problem, <u>d</u>ynamic <u>t</u>ime <u>w</u>arping (DTW) [25] was used to adjust the time-axis of the data points in *T* to match the empirical data points in *S* (Supplemental Fig. 1). DTW tends to have the singularity problem by matching the signal of a short time window to that of a long time window [26], so a constraint was introduced such that the ratio between any two time periods matched by DTW should be between 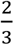 and 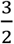. That is to say, 6 data points in *T* can be matched with at least 4 data points and at most 9 data points in *S*. The squared error was used to measure the distance between two data points *s_i_* and *t_i_*, i.e., *d*(*s_i_, t_i_*) = (*s_i_ – t_i_*)^2^.

We fill out a 2-dimensional (*m* + 1) × (*m* + 1) table *D*, in which *D*[*i, j*] stores the minimum distance between *S*[1,*i*] and *T*[1,*j*] after time warping. The recurrence function for computing *D*[*i,j*] is shown Step 4 in Supplemental Fig. 1. Because at least 2 data points in *S* are needed to match 3 data points in *T* and *vice versa*, the singularity problem is solved. The time complexity of the algorithm is *O*(*m*^2^).

### 2.9 Consensus nanospectra

To reduce the noise in nanospectra, a consensus spectrum of a peptide was formed by combining all nanospectra of the peptide. Accordingly, if *S*_1_,*S*_2_,…,*S_n_* are the nanospectra of a peptide after orientation correction and *S_i_*[*j*] is the current blockade signal for the *j*^th^ point in *S_i_* for 1 ≤ *i* ≤ *n* and 1 ≤ *j* ≤ *m*, then the consensus spectrum *S* was formed by taking the average current blockade signals of the nanospectra. That is to say, the consensus signal 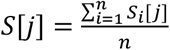 for 1 ≤ *j* ≤ *m*. The nanospectrum *S* is called the *average consensus nanospectrum* of the peptide.

One limitation of the average consensus approach was that it failed to consider the variance in the velocity with which AAs pass the sub-nanopore. The relative dwell time of an AA in a peptide molecule is the ratio between the AA dwell times and the whole molecule. The relative dwell times in nanospectra for the same AA in the peptide could be different. Owing to this variance, the current blockade signals *S*_1_[*j*],*S*_2_[*j*],…,*S_n_*[*j*] for the same position *j* could originate from different AAs and so the average current blockade signal may be an inaccurate consensus of the nanospectra.

Similar to multiple sequence alignment [27], a progressive method was used to improve the quality of average consensus nanospectra with high-quality empirical nanospectra (Supplemental Fig. 2). According to this algorithm, DTW was used to align each empirical nanospectrum with the *average consensus nanospectrum*, and then each was ranked in the increasing order of the distance. The top *t* empirical nanospectra (*t* = 50 in this analysis) were chosen to update the consensus. The best empirical nanospectrum was first aligned with the average consensus nanospectrum, and the average consensus nanospectrum was then updated by forming a weighted average with the best empirical nanospectrum. This step was repeated for the top *t* nanospectra. Specifically, to update the consensus using the *i*th empirical spectrum, the weight for the consensus was *u+i-1* and that for the highly ranked empirical spectrum was 1, where *u* is the weight for the original consensus (*u* = 30 in the experiments). The updated consensus nanospectrum is referred to as *the alignment consensus nanospectrum* of the peptide.

The functions for reading ABF files were implemented in MATLAB, whereas all the other functions were coded in Python. All the data processing was performed on a computer with an Intel Core i7-6700 3.4 GHz CPU and 16 GB memory.

## 3. Results

### 3.1 Sub-nanopore fabrication and characterization

A sub-nanopore sputtered through a thin, nominally 5 nm thick, a-*Si* membrane was used to analyze the peptides. The thickness was important because it affected the field distribution in the pore and therefore the resolution of a read. A pore was sputtered in the window through the a-*Si* membrane using a tightly focused, high energy (300 keV) electron beam formed in either an FEI Titan or Themis Z STEM. Subsequently, the pore was visualized *in situ* with TEM immediately after sputtering to reveal a 1.0 ×1.5 nm^2^-cross-section at the waist defined by the shot noise (Fig. 1a). However, the pore topography was likely affected, not only by electron-beam sputtering but also by oxidation in the ambient. This is likely because after exposure to the ambient for 1–3 days, the same pore was re-acquired and the topography visualized with HAADF using an aberration-corrected (Themis Z) STEM (Fig. 1b) to reveal a smaller lumen [17]. Based on images like this, the pore topography was bi-conical with a steep cone angle > 7.4° that broadened to 16° near the orifice with an irregular waist 0.65 nm × 0.87 nm in cross-section.

**FIGURE 1.**
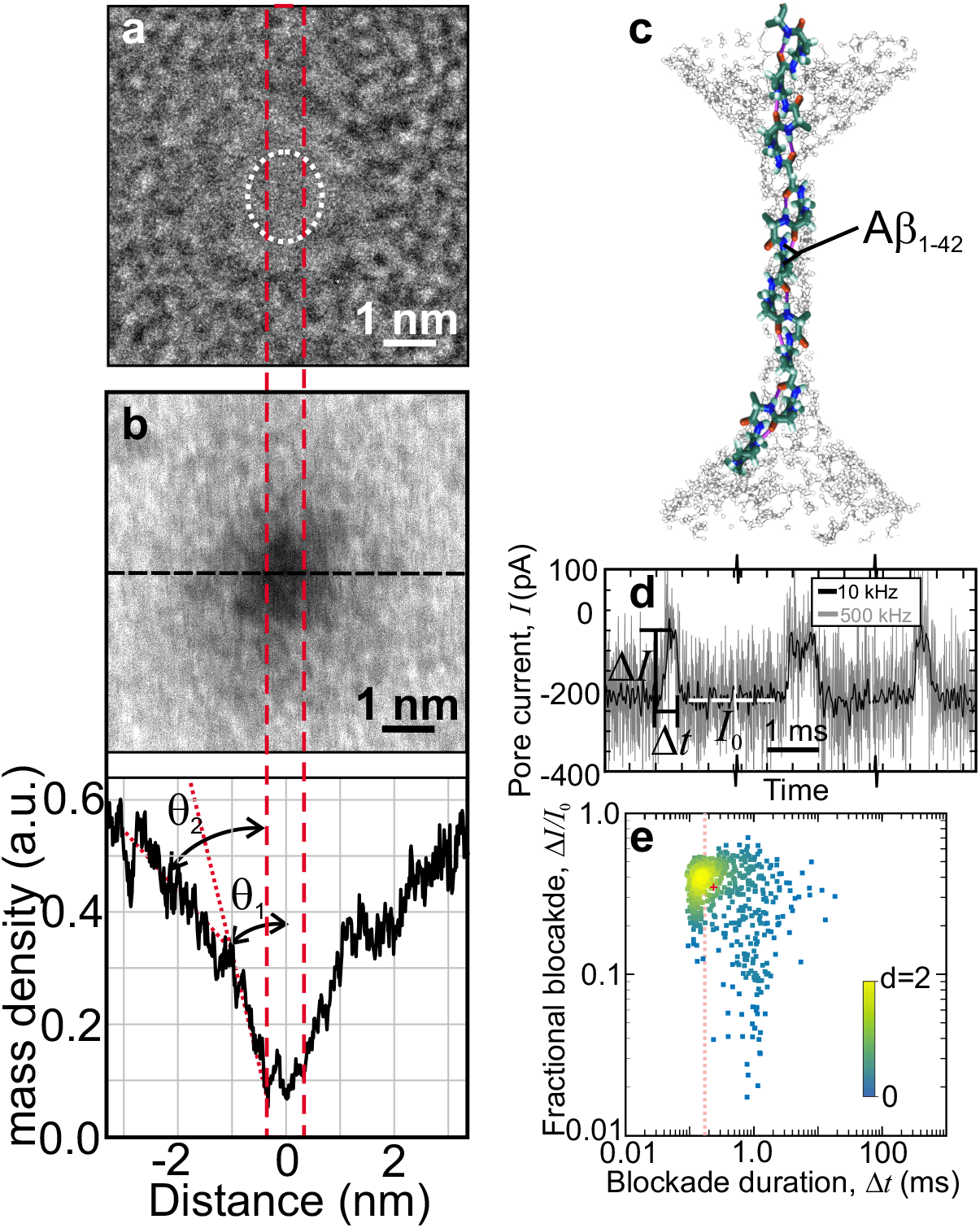
Improved read resolution and fidelity using a sub-nanopore through a thin laminated a-*Si* membrane. **(a)** A TEM image is shown *in vacuo* of a nanopore immediately after sputtering through a nominally 5 nm thick a-*Si* membrane. The cross-section of the pristine pore was estimated from the shot noise associated with electron transmission through the pore to be about 1.0 × 1.5 nm^2^ (dotted circle). **(b, top)** An HAADF-STEM image, acquired with an aberration-corrected microscope is shown of the pore in (a) after exposure to the ambient. **(b, bottom)** The profile of the mass-density under the probe beam is shown taken along the dashed (horizontal) line in (d, top). The cross-section shrunk to about 0.65 nm × 0.87 nm, indicative of the growth of a native oxide in the pore waist. **(c)** A schematic representation is shown depicting a translocation of Aβ_1-42_ impelled by an electric force through a sub-nanopore. The actual pore is ghosted in the figure; only the peptide is represented. **(d)** Current traces (negative raw current) are shown that illustrate the distribution of the duration of the blockade currents associated with translocations of single molecules of Aβ_1-42_ through a sub-nanopore spanning an a-*Si* membrane at 0.6 V. The pore current was amplified over a >75 kHz bandwidth and sampled at 500 kHz (gray line) to detect each residue in the peptide in a Δ*t* = 170 μs blockade. Another version of the same data, filtered with a 10 kHz eight-pole Bessel filter (black line), is also shown. The definition of the blockade current, Δ*I*, the blockade duration, Δ*t*, and the open pore current, *I*_0_, are indicated. Higher current (negative raw current) values correspond to larger blockade currents. **(e)** A heat map is shown that illustrates the distribution of fractional blockades relative to the open pore current (Δ*I/I*_0_) versus the blockade duration (Δ*t*) associated with single denatured Aβ_1-42_ molecules translocating through a sub-nanopore acquired at 0.6 V. The red dotted line shows the position of 170 μs.

If the bi-conical topography focussed the electric field to a sub-nanometer extent near the waist then it followed that a blockade mainly measured the occluding volume due to the AAs in the waist (Fig. 1c). So, if only a few acids occupied the waist at a time, it was reasoned that the blockade current would mainly measure the volume associated with those residues. Likewise, as it has been shown empirically that the small size of a sub-nanopore knocks-down the mobility of de-hydrated ions [5], so it should also affect the acid mobility in the same way. Doubtless other AAs outside the waist would still contribute at least marginally to the blockade current and the mobility in the pore.

Heat (85°C), SDS, and BME were used to denature the peptide and maintain it. SDS is an anionic detergent that works, in combination with heat and reducing agents like BME, to impart a nearly uniform negative charge to the protein that stabilizes denaturation. Although the exact structure of the aggregate formed by SDS and protein remains unsolved, a “rod-like” model was adopted in which the SDS molecules form a shell along the length of the protein backbone [28]. The resulting uniform charge on the protein was supposed to facilitate electrical control of the translocation kinetics. Due to its size, however, it is unlikely that the SDS remained bound to the protein as the aggregate was forced through the sub-nanopore by an applied electric field. Rather, it was likely cleaved from the protein by the steric constraints imposed by the pore topography above the waist [3].

### 3.2 Measurements of the blockade current

Measurements of fluctuations in the blockade current through a sub-nanopore were used to analyze the acid sequence of two synthesized peptides: a 42-residue (human) amyloid-β (Aβ)-protein fragment Aβ_1-42_ and a scrambled variant SAβ_1-42_ of it (Methods) and have been reported in [17]. The blockade is defined as the difference between the open sub-nanopore current *I*_0_ and the current *I* in the peptide translocation, that is, Δ*I* = *I*_0_ – *I*. When a nearly pH-neutral (pH 6.6 ± 0.1) solution containing denatured Aβ_1-42_ or SAβ_1-42_ peptides was introduced on the *cis*-side of a sub-nanopore with a voltage of 0.40-0.6 V applied across the membrane, blockades were observed almost immediately (Fig. 1d). The blockades were attributed to the translocation of rodlike single peptides across the membrane through the sub-nanopore (Fig. 1c). To account for the rapidity of the translocation, the electrical signal was amplified over a 75 kHz bandwidth and sampled at 500 kS/s. Accordingly, the signal was obscured by electrical noise.

Clusters of blockades were selected in a range demarcated by the Nyquist sampling rate corresponding to at least 0.5 samples per AA (with a blockade duration Δ*t* > 42 μs for Aβ_1-42_ and SAβ_1-42_ amplified with a 75-100 kHz bandwidth, and then sampled at 500 kS/s). To facilitate comparisons, the selected blockades of Aβ_1-42_ were classified by the duration of the blockade (Δ*t*) and the *fractional blockade*, which is the ratio between in the blockade current and the open sub-nanopore current (Δ*I*/*I*_0_). The aggregate data was then represented by normalized heat maps of the <u>p</u>robability <u>d</u>ensity <u>f</u>unctions (PDFs) reflecting the number and distribution of blockades (Fig. 1e). Almost all the blockades have a duration longer than 42 μs, whereas about half had a duration >170 μs (Fig. 1e). Blockades that were too short in duration could not realistically inform on all the residues with the limited bandwidth of the amplifier and the 500 kS/sec sampling rate. On the other hand, blockades that were too long would likely muddle the interpretation of the signal because of (slip-stick) translocation kinetics [3]. In data preprocessing, blockades with a long duration were still included because they can provide some information of AAs, and all blockades with a duration < 170 μs were removed, resulting in 475 and 2,000 nanospectra for Aβ_1-42_ and S Aβ_1-42_, respectively (Methods).

### 3.3 Consensus nanospectra

The orientations of nanospectra were determined using the PCCs between empirical nanospectra and theoretical ones generated from the 1AAV model. Of the 475 Aβ_1-42_ nanospectra, the orientations of 268 were forward and 207 were backward. Of the 2,000 SAβ_1-42_ nanospectra, 950 were forward and 1,050 were backward. Many empirical spectra have a small difference between the PCCs of the original nanospectrum and the flipped one, making it challenging to confidently determine their orientations (Supplemental Fig. 3).

An *average consensus nanospectrum* of the peptide was formed to recover reproducible fluctuations in the blockade signal from irreproducible noise. The average consensus nanospectra were aligned with the corresponding theoretical nanospectra (1AAV) using DTW. It was compelling that the amplitude fluctuations in the average consensus nanospectra (Figs. 2a,b; orange lines) were highly correlated to the theoretical nanospectra (Figs. 2a,b; blue lines). Strikingly, the amplitude of the fluctuations tracked the AA volumes ascribed to the primary structure of Aβ_1-42_ with PCC = 0.896 (Fig. 2a). A sub-nanopore assay of SAβ_1-42_, consisting of a different sequence of the same residues produced conspicuous differences in the fluctuation pattern (Fig. 2b), however, and was correlated (PCC = 0.880) to the corresponding 1AAV model for the scrambled sequence.

**FIGURE 2.**
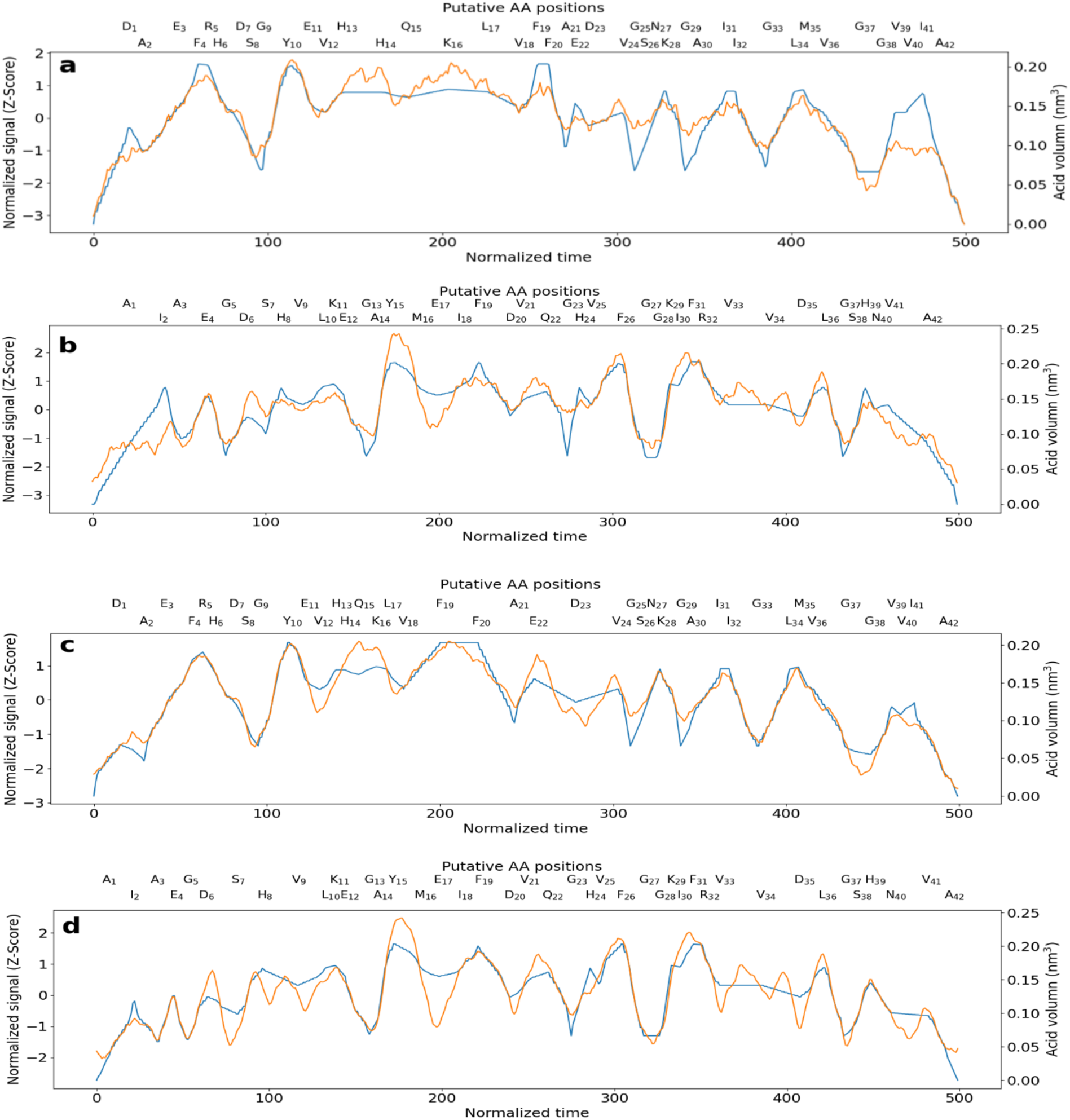
**(a)** A plot of a 475-blockade average consensus nanospectrum acquired at 0.6 V by forcing denatured Aβ_1-42_ through a sub-nanopore is shown versus normalized duration (orange line). Aligned with the empirical data is the corresponding 1AAV model (blue line) using DTW. The blockade current was correlated (PCC = 0.896) with the corresponding volume model. **(b)** A plot of a 2000-blockade average consensus nanospectrum acquired at 0.6 V by forcing denatured SAβ_1-42_ through a sub-nanopore is shown versus normalized duration (orange line). Aligned with the empirical data is the corresponding 1AAV mode (blue line) with DTW. The empirical consensus was correlated (PCC = 0.880) with the corresponding 1AAV model. **(c)** The alignment consensus nanospectrum (orange line) of Aβ_1-42_ is aligned with the 1AAV-P model (blue line) with PCC = 0.954. **(d)** The alignment consensus nanospectrum (orange line) of SAβ_1-42_ is aligned with the 1AAV-P model (blue line) with PCC = 0.903.

The *average consensus nanospectra* for Aβ_1-42_ and SAβ_1-42_ were further improved by using the progressive alignment method with the parameter *u* set to 30 (Methods). The PCCs for the *alignment consensus nanospectra* and the theoretical nanospectra (1AAV) were 0.919 and 0.876 for Aβ_1-42_ and SAβ_1-42_, respectively (Supplemental Fig. 4). The progressive alignment method increased the quality of the Aβ_1-42_ consensus nanospectrum but lowered slightly the quality of the SAβ_1-42_ consensus nanospectrum. It is likely that the top empirical nanospectra of Aβ_1-42_ forming the consensus might be of higher quality than those of SAβ_1-42_, so they could improve the consensus.

The correlations that developed between the consensus nanospectra and the corresponding volume models were important for two reasons. The fluctuations translated to reads with (nearly) single residue resolution, which could facilitate calling AAs as it alleviates the analytical and computational burden associated with ferreting out the identity of multiple monomers producing a fluctuation in a blockade. Second, it was also important because the fidelity proves that the signal-noise ratio can be improved with a reduction of the parasitic capacitance and with enough signal averaging, even with a high sampling frequency and no filtering. The correlation between the empirical consensuses and the corresponding volume models used for AA calls was still imperfect.

### 3.4 Prediction of blockade currents

Seeking further refinement of the model, the alignment consensus and theoretical (1AAV) nanospectra of Aβ_1-42_ were normalized using zero mean and unit variance to form the Z-score, and then 42 data points were extracted from the alignment between the consensus and theoretical nanospectra. Each data point corresponded to an AA in the peptide. Likewise, 42 data points were extracted from the SAβ_1-42_ alignment consensus nanospectrum. Then linear regression was used to predict blockade signals with six encoding methods: 1AAV, 3AAV, AAG, 1AAV-P, 3AAV-P, and AAG-P (see Methods). Prediction accuracy was evaluated using 2-fold cross-validation: first, the training data were the Aβ_1-42_ data points and the validation data were the SAβ_1-14_ data points, and then the training and validation data sets were swapped. The error function was the <u>m</u>ean <u>s</u>quared <u>e</u>rror (MSE).

The methods with the position feature outperformed those without the feature, showing that the positions of AAs affect their current blockade signals, especially for those near the N- or C-terminus (Table 1). The AAs near the N- or C-terminus tend to have lower blockade signals than those in the middle. The 1AAV-P method obtained the best validation error. 3AAV-P and AAG-P reported better training errors than 1AAV-P, but their validation errors were worse than 1AAV-P, showing that they might have an overfitting problem due to the limited size of the training data. We also tested support vector machine (SVM) regression and random forest regression, but their performance was not as good as linear regression.

**TABLE 1.** Comparison of six encoding methods for predicting blockade signals

|  | 1AAV | 3AAV | AAG | 1AAV-P | 3AAV-P | AAG-P |
| --- | --- | --- | --- | --- | --- | --- |
| Training error (MSE) | 0.256 | 0.229 | 0.221 | 0.193 | 0.186 | **0.169** |
| Validation error (MSE) | 0.256 | 0.259 | 0.289 | **0.211** | 0.223 | 0.241 |

The AA positions and volumes were incorporated (1AAV-P) into revised estimates for the theoretical nanospectra of Aβ_1-42_ and SAβ_1-42_. When the position feature *x_P_* of an AA is less than 5, the volume of the AA was adjusted by −11.3(5 – *x_P_*). The parameter −11.3 was estimated based on the coefficients reported by linear regression. With the adjusted volumes, the PCCs between theoretical and consensus nanospectra were improved to 0.954 and 0.903 for Aβ_1-42_ and SAβ_1-42_, respectively (Figs. 2c,d).

### 3.5 Statistical significance of nanospectral identifications

Finally, 10,000 random peptides 42 acids long and their corresponding theoretical nanospectra were generated using the 1AAV-P method. Subsequently, DTW was used to align the theoretical spectra and the *alignment consensus nanospectrum* of Aβ_1-42_. The average and best PCCs of the random peptides were 0.839 and 0.959, respectively (Supplemental Fig. 5). Based on the PCCs of the random peptides, the estimated *p*-value of the match between the theoretical and *alignment consensus nanospectra* of Aβ_1-42_ was about 0.0003, which is statistically significant enough for peptide identification when the database is not very large.

## 4. Conclusions and discussion

Various computational methods for signal processing, blockade current prediction, and identification of nanospectra using Aβ_1-42_ and SAβ_1-42_ peptides have been scrutinized for protein sequencing and identification. Since raw nanospectra are noisy, an indispensable pre-processing step is to use average nanospectra and alignment to obtain a high-quality consensus nanospectrum. Progressive alignment between the average consensus and top raw nanospectra could further improve the consensus of Aβ_1-42_, but not SAβ_1-42_. Apparently, the performance of the alignment method depends on the quality of raw nanospectra.

Six methods for predicting blockade signals of AAs were tested and benchmarked. By adding the positional information into blockade signal prediction, the PCCs between theoretical and empirical nanospectra were improved. Because only 84 data points were used for training and validation, only the 1AAV method showed similar accuracy in training and validation. The 3AAV and AAG methods obtained small prediction errors in the training data, but their validation errors were large. These methods have the potential to improve prediction accuracy, but more training data are needed to address the overfitting problem.

The estimated *p*-value of the match between the theoretical and alignment consensus nanospectra of Aβ_1-42_ was 0.0003. Thus, peptides can be identified unambiguously using the nanospectra from a database of thousands of peptides, showing the potential of sub-nanopore sequencing to identify peptides from a peptide mixture.

There are many computational problems in nanospectral data analysis that have not been well studied. Nanospectral clustering is an important pre-processing step for analyzing nanospectra of peptide mixtures. Predicting the peptide length of nanospectra is needed to identify truncated proteoforms. There are still no software tools for these problems. Accurate theoretical nanospectra can significantly increase the statistical significance of identifications in database search. So further improvement in the accuracy is needed for predicting theoretical nanospectra of peptides and those with PTMs—molecular dynamics simulations may be useful in this endeavor. *De novo* peptide sequencing from nanospectra is a challenging problem with high impact. A large nanospectral data set is also needed for training machine learning models and test the performance of nanospectral data analysis methods.

## Acknowledgments

This research was supported by a grant from the Open Philanthropy Project and partially supported by the Keough-Hesburgh Professorship.

