## Supplementary Material for "Calling the Amino Acid Sequence of a Protein/Peptide from the Nanospectrum Produced by a Sub-nanometer Diameter Pore"

**Input:** An experimental nanospectrum $S = s_1 s_2 \ldots s_m$ and a theoretical nanospectrum $T = t_1 t_2 \ldots t_m$

**Output:** An alignment between $S$ and $T$

1. Initialize an $(m+1) \times (m+1)$ table $D$ by setting $D[0,0] = 0$, $D[1,1] = d(s_1, t_1)$ and other cells to $\infty$. We also assume $D[i,j] = \infty$ when $i < 0$ or $j < 0$.
2. **For** $i = 2$ to $m$ **do**
3. **For** $j = 2$ to $m$ **do**
4. $$D(i,j) = \min \begin{cases} D(i-2, j-2) + d(s_{i-1}, t_{j-1}) + d(s_i, t_j) \\ D(i-2, j-3) + d(S[i-1,i], T[j-2,j]) \\ D(i-3, j-2) + d(S[i-2,i], T[j-1,j]) \end{cases}$$
5. Use backtracking to find a best alignment between $S$ and $T$.

**SUPPLEMENTARY FIGURE 1**. Dynamic time warping algorithm with a constraint. The distance between two data points $S[i-1,i]$ and three data points $T[j-2,j]$ is defined as $d(S[i-1,i], T[j-2,j]) = d(s_{i-1}, t_{j-2}) + d(s_i, t_j)$. The distance between $S[i-2,i]$ and $T[j-1,j]$ is defined as $d(S[i-2,i], T[j-1,j]) = d(s_{i-2}, t_{j-1}) + d(s_{i-1}, t_j) + d(s_i, t_j)$.

**Input:** An average consensus nanospectrum $C$, a list of experimental nanospectra $S_1, S_2, \ldots, S_n$ in the increasing order of their distances with $C$, and parameter $u$.

**Output:** An improved consensus nanospectrum

1. **For** $i = 1$ to $m$ **do**
2. Use DTW to align $C$ and $S_i$. The nanospectrum $S_i$ after time warping is represented by $S_i'$.
4. Update $C$ using the weighted average of $C$ and $S_i'$. The weights of $C$ and $S_i'$ are $u + i - 1$ and 1, respectively.
4. Return $C$.

**SUPPLEMENTARY FIGURE 2**. Algorithm for improving the average consensus nanospectrum by alignment.

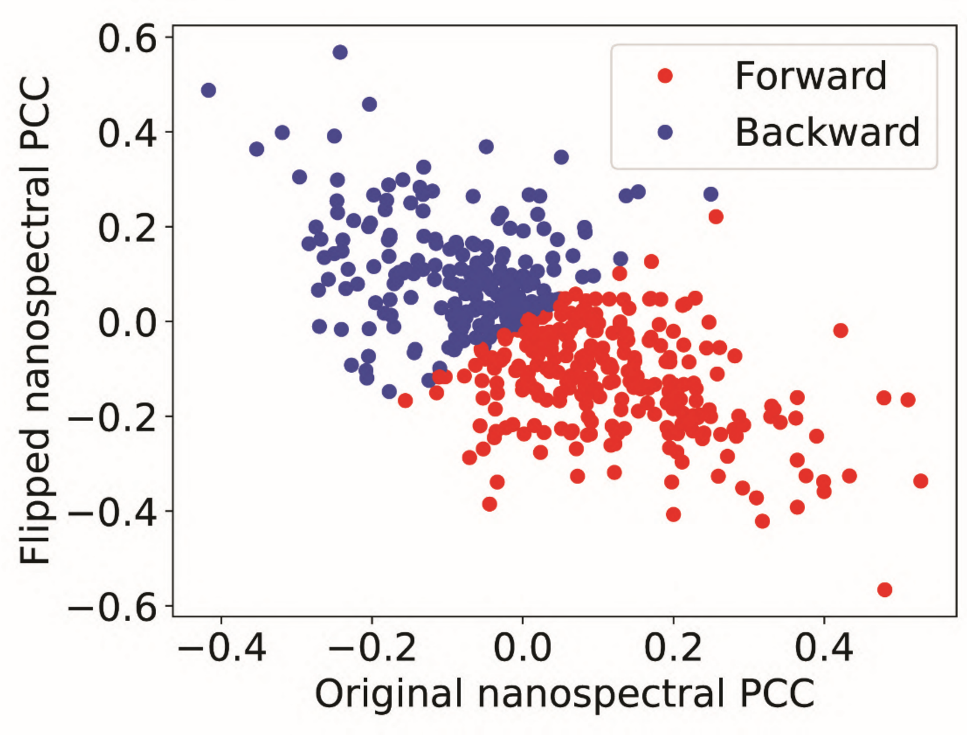


**SUPPLEMENTARY FIGURE 3**. PCCs of the empirical nanospectra and flipped empirical nanospectra of A$\beta_{1\text{-}42}$ compared with the theoretical nanospectrum generated using 1AAV model and linear interpolation.

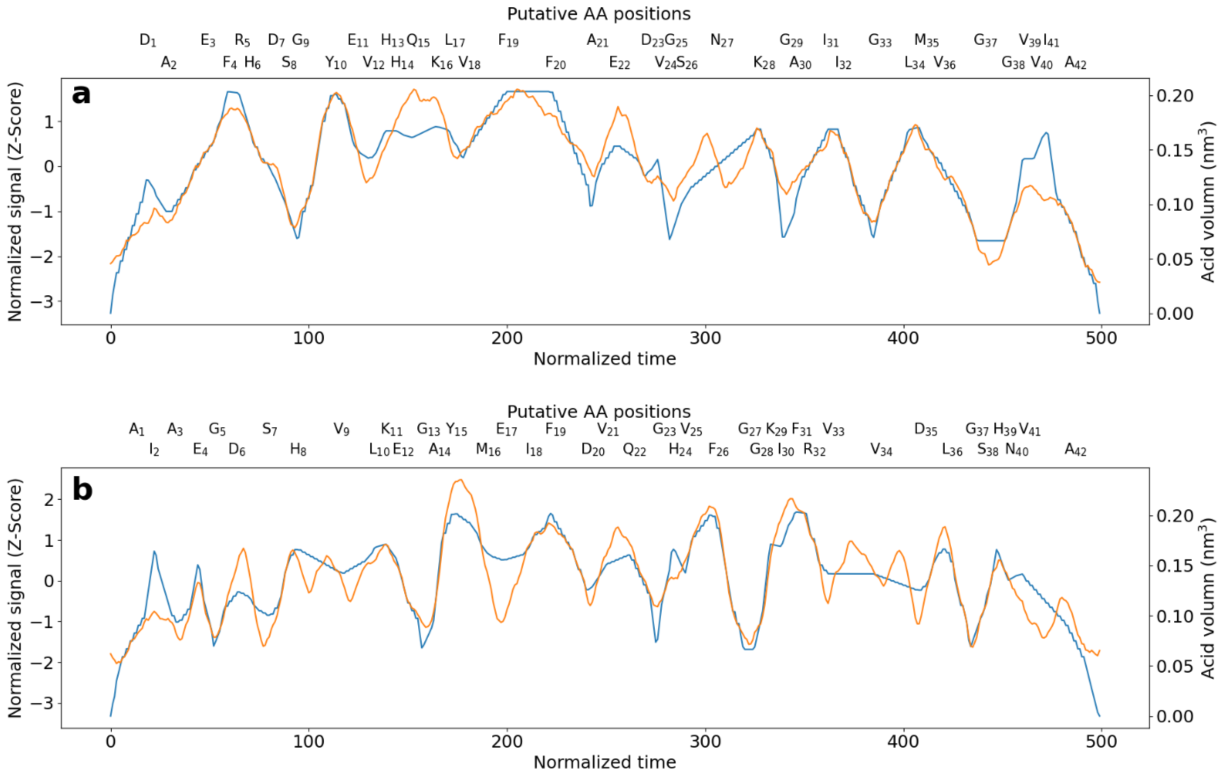


**SUPPLEMENTARY FIGURE 4. (a)** A plot of a 475-blockade alignment consensus nanospectrum of $A\beta_{1-42}$ is shown versus normalized duration (orange line). Aligned with the empirical data is the corresponding 1AAV model (blue line) using DTW. The alignment consensus was correlated (PCC = 0.919) with the corresponding volume model. **(b)** A plot of a 2000-blockade alignment consensus nanospectrum of $SA\beta_{1-42}$ is shown versus normalized duration (orange line). Aligned with the empirical data is the corresponding 1AAV mode (blue line) with DTW. The empirical alignment consensus was correlated (PCC = 0.876) with the corresponding 1AAV model.

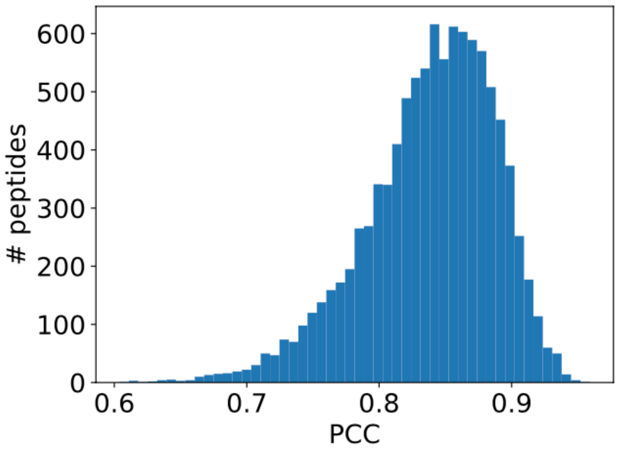


**SUPPLEMENTARY FIGURE 5.** Distribution of the PCCs between the alignment consensus nanospectra of A$\beta_{1-42}$ and the theoretical nanospectra of 10,000 random peptides after DTW.

1. Rigo, E., et al., *Measurements of the size and correlations between ions using an electrolytic point contact.* Nat Commun, 2019. **10**(1): p. 2382.
